# Diagnosis and analysis of unexplained cases of childhood encephalitis in Australia using metagenomic next-generation sequencing

**DOI:** 10.1101/2021.05.10.443367

**Authors:** Ci-Xiu Li, Rebecca Burrell, Russell C Dale, Alison Kesson, Christopher C Blyth, Julia E Clark, Nigel Crawford, Cheryl A. Jones, Philip N. Britton, Edward C. Holmes, on behalf of the Australian Childhood Encephalitis study investigators

## Abstract

Encephalitis is most often caused by a variety of infectious agents, the identity of which is commonly determined through diagnostic tests utilising cerebrospinal fluid (CSF). Immune-mediated disorders are also a differential in encephalitis cases. We investigated the clinical characteristics and potential aetiological agents of unexplained encephalitis through metagenomic next-generation sequencing of residual clinical samples of multiple tissue types and independent clinical review. A total of 43 specimens, from both sterile and non-sterile sites, were collected from 18 encephalitis cases with no cause identified by the Australian Childhood Encephalitis study. Samples were subjected to total RNA sequencing to determine the presence and abundance of potential pathogens, to reveal mixed infections, pathogen genotypes, and epidemiological origins, and to describe the possible aetiologies of unexplained encephalitis. From this, we identified five RNA and two DNA viruses associated with human infection from both non-sterile (nasopharyngeal aspirates, nose/throat swabs, urine, stool rectal swab) and sterile (cerebrospinal fluid, blood) sites. These comprised two human rhinoviruses, two human seasonal coronaviruses, two polyomaviruses and one picobirnavirus. With the exception of picobirnavirus all have been previously associated with respiratory disease. Human rhinovirus and seasonal coronaviruses may be responsible for five of the encephalitis cases reported here. Immune-mediated encephalitis was considered clinically likely in six cases and RNA sequencing did not identify a possible pathogen in these cases. The aetiology remained unknown in nine cases. Our study emphasises the importance of respiratory viruses in the aetiology of unexplained child encephalitis and suggests that the routine inclusion of non-CNS sampling in encephalitis clinical guidelines/protocols could improve the diagnostic yield.

**Author Summary:** Encephalitis is caused by both infectious agents and auto-immune disorders. However, the aetiological agents, including viruses, remain unknown in around half the cases of encephalitis in many cohorts. Importantly, diagnostic tests are usually based on the analysis of cerebrospinal fluid which may limit their utility. We used a combination of meta-transcriptomic sequencing and independent clinical review to identify the potential causative pathogens in cases of unexplained childhood encephalitis. Accordingly, we identified seven viruses associated with both sterile and non-sterile sampling sites. Human rhinovirus and seasonal coronaviruses were considered as most likely responsible for five of the 18 encephalitis cases studied, while immune-mediated encephalitis was considered the cause in six cases, and we were unable to determine the aetiology in nine cases. Overall, we demonstrate the role of respiratory viruses as a cause of unexplained encephalitis and that sampling sites other than cerebrospinal fluid is of diagnostic value.

## Introduction

Encephalitis, defined as inflammation of the brain tissue, is caused by a broad range of infectious agents, including bacteria, viruses, fungi and parasites, as well as a number of auto-immune disorders [1, 2]. However, in 40% to 60% of cases in many cohorts the aetiological agents remain unknown [3, 4]. Investigation is challenging because clinical presentation may be atypical and the differential diagnosis may include unusual and opportunistic pathogens – especially if the patient is immunosuppressed - or a non-infective cause. The conventional means of infectious encephalitis diagnosis are specific PCR assays on central nervous system samples including cerebrospinal fluid (CSF) and brain biopsy, as well as immunohistochemical and serological assays [4]. Failure to detect a causative agent by such routine laboratory diagnostics may reflect a lack of diagnostic tests for rare, novel or divergent pathogens, limited volume of CNS samples, and overlapping clinical presentation caused by infectious and non-infectious processes [5].

The incidence of all-cause childhood encephalitis (including infection-associated encephalopathy) is estimated to be between 3.8 and 5.0 per 100,000 population aged ≤14 years [6]. The leading causes are picornaviruses (enteroviruses and parechoviruses), herpes simplex viruses 1 and 2, influenza (infection associated encephalopathy), bacterial meningo-encephalitis and immune-mediated causes such as acute disseminated encephalo-myelitis (ADEM) and anti-N-methyl-D-aspartate receptor (NMDAr) encephalitis [7]. The burden of disease is considerable, with 49% admitted to ICU, a case-fatality rate of 5%, and 27% of cases showing neurodisability at discharge from hospital [7].

Metagenomic next-generation sequencing (mNGS) has successfully identified a broad range of infectious agents in a range of clinical syndromes [8–10] and is gradually being established as a powerful and reliable diagnostic platform [8]. Indeed, mNGS has identified an increasing number of novel or unexpected pathogens associated with encephalitis [11]. Total RNA sequencing – meta-transcriptomics – may be especially powerful as it provides a simple way to characterise all the actively transcribing microbes in a sample and estimate their abundance [12–14]. Not only does total RNA sequencing identify the RNA viruses present in a sample, but also those DNA viruses, bacteria, parasites and fungi that are actively transcribing RNA [15].

The Australian Childhood Encephalitis (ACE) study has comprehensively identified, collected data, and reviewed cases of this severe syndrome nationally through active sentinel hospital surveillance since 2013, and requested banking of salvaged laboratory biospecimens from cases [7]. Herein, we describe the use of total RNA sequencing on samples of differing tissue types obtained from 18 cases of childhood encephalitis categorised as having unknown cause. Notably, several common respiratory viruses, including human seasonal coronaviruses and rhinoviruses, were identified in samples outside of the CNS and potentially caused the disease process in these cases.

## Results

### Clinical evaluation

A total of 18 cases, representing patients from four Australian states (Figure 1A), were chosen for study. These cases had previously been reviewed and confirmed to have encephalitis using the Brighton criteria and International Encephalitis Consortium (IEC) definitions [7], but with unknown causality (Table 1). The sex ratio of patients was 1.57:1 male to female, with ages ranging from neonates to twelve years (average, 4.5 years) such that only pre-adolescent children were considered (Table 1).

**Fig. 1.**
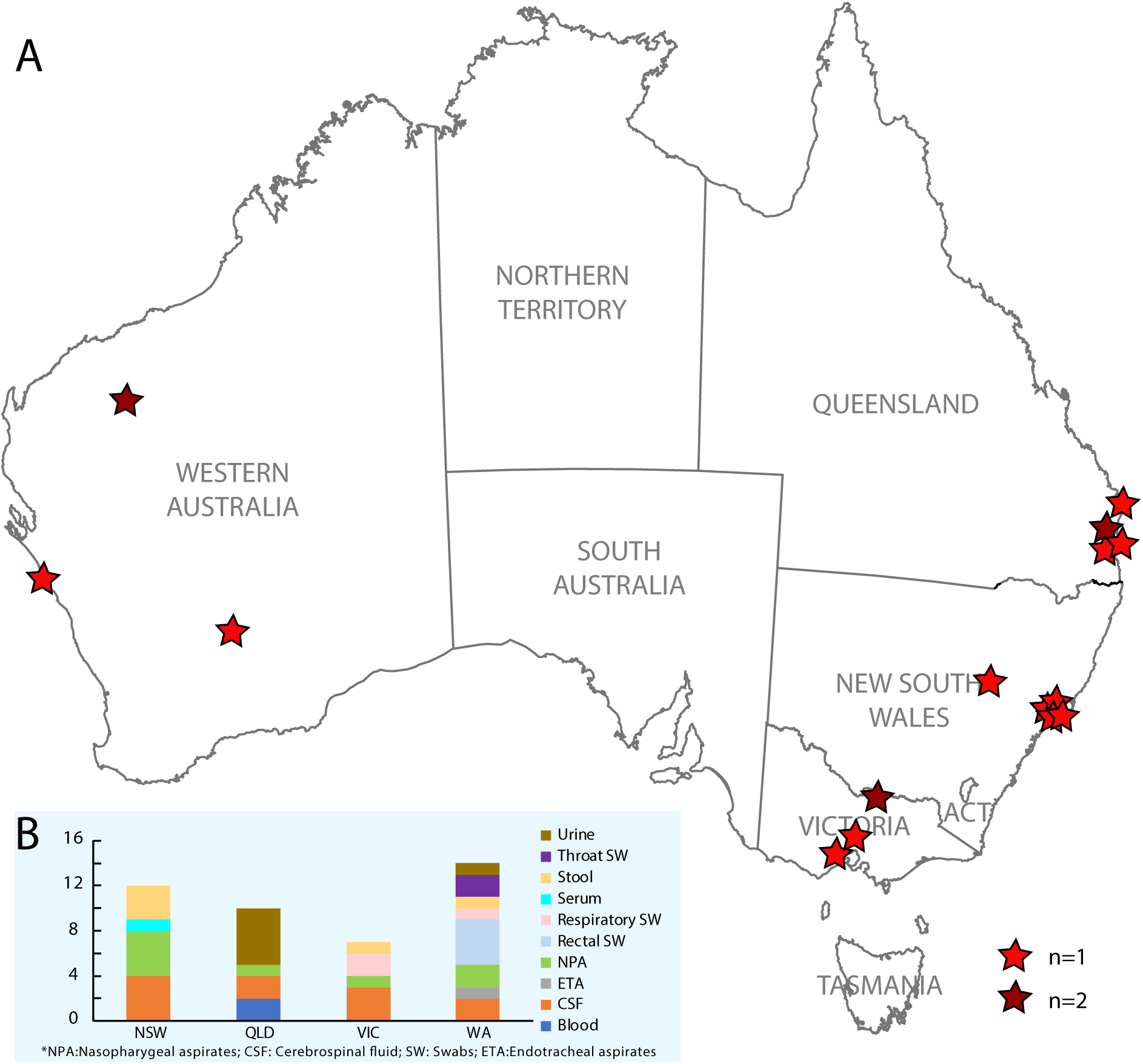
Geographic context of the study. (A) Maps showing the residential postal codes within Australia of patients with clinical diagnosis as encephalitis reported at the Children’s Hospital at Westmead (CHW) between March 2014 and September 2016. Red star indicates one case and dark red star indicates two cases. (B) The bar chart showing the diversity of sample types collected in four states (NSW, QLD, VIC and WA) in Australia, coloured by sample types.

**Table 1.** Demographic and clinical characteristics of the 18 patients studied here.

| Case No. | Sex/Age | Respiratory symptoms | Gastro-intestinal symptoms | CNS Inflammation |  |  |  |  | Reduced Responsiveness |  |  |  | *Focal neurological signs | Brighton criteria | International encephalitis consortium criteria |
| --- | --- | --- | --- | --- | --- | --- | --- | --- | --- | --- | --- | --- | --- | --- | --- |
|  |  |  |  | Fever | CSF pleocytosis | CSF WCC | MRI changes | EEG changes | Absent environment | Absent eye | Decreased arousability | Seizure LOC |  |  |  |
| 1 | F/3.2mo |  | Y | Y | Y | 11 | Y | Y | Y | Y | Y | Y | Y | Level 2 | Probable |
| 2 | M/7.1yo |  | Y | Y | Y | 501 |  | N/A | Y |  |  |  | Y | Level 2 | Possible |
| 3<sup>#</sup> | M/1.1yo |  | Y | Y |  | 3 | Y |  | Y | Y | Y | Y | Y | Level 2 | Probable |
| 4 | M/7.4yo |  | Y | Y |  | 2 | N/A | Y | Y |  | Y |  |  | Level 2 | Possible |
| 5 | M/9.5yo |  |  | Y |  | 0 |  | Y | Y | Y | Y |  | Y | Level 2 | Possible |
| 6 | M/2.6yo |  |  | Y |  | 1 |  | N/A | Y | Y | Y |  | Y | Level 3 | Possible |
| 7 | F/1.1yo |  | Y |  | Y | 21 | Y | Y | Y | Y | Y | Y | Y | Level 2 | Probable |
| 8<sup>†</sup> | M/8.5yo |  |  | Y | Y | 29 | Y | N/A | Y | Y | Y |  | Y | Level 2 | Probable |
| 9 | F/3.8wks |  |  | Y | Y | 65 | Y | Y | Y |  | Y |  | Y | Level 2 | Possible |
| 10<sup>†</sup> | M/9.9yo |  | Y |  | Y | 29 | Y | N/A |  |  | Y |  |  | Level 2 | Probable |
| 11 | M/3mo |  |  |  | Y | 72 | Y | N/A | N/A | Y |  |  | Y | Level 3 | Probable |
| 12<sup>†</sup> | M/3.6yo |  | Y |  |  | 0 | Y | N/A |  |  |  |  | Y | Level 3 | Probable |
| 13 | F/3.4yo | Y | Y | Y | Y | 21 | Y |  | Y | Y | Y | Y | Y | Level 2 | Probable |
| 14 | M/12.6yo | Y | Y |  | Y | 66 |  | N/A | Y | Y | Y | Y | Y | Level 2 | Possible |
| 15 | M/8.1yo |  |  | Y |  | 6 | Y | Y | Y | Y | Y |  | Y | Level 2 | Probable |
| 16 | F/8.6mo |  |  |  | Y | 31 | Y | Y | Y | Y | Y | Y | Y | Level 2 | Probable |
| 17<sup>##</sup> | F/3.3yo |  |  | Y | N/A | N/A | N/A |  |  |  | Y | Y | Y | Level 3 | Probable |
| 18 | F/7.8yo |  |  | Y | Y | 7 | Y | Y | Y | Y | Y | Y | Y | Level 2 | Probable |

Y = Yes, N/A = Not Done. All cases had an initial expert panel diagnosis of Unknown Encephalitis. \* Y is indicative of cases with one or more of the following: focal seizure, cranial nerve abnormality, visual impairment, motor weakness, altered deep tendon reflexes, cerebellar dysfunction, abnormal movement, or any other focal neurological signs. Respiratory symptoms included cough/coryza and increased work of breath requiring mechanical ventilation for both cases, gastrointestinal symptoms include diarrhoea and/or vomiting.
<sup>#</sup>This patient was with severe combined immunodeficiency (SCID) and the outcome was death.
<sup>##</sup>This patient was reported with respiratory illness in the previous 2 weeks before admission.
<sup>†</sup>These patients were transferred to intensive care unit (ICU) within 24 hours of admission.

Of the 18 unknown encephalitis patients, abnormal neuroimaging (MRI/CT)) was found in 14/18, and EEG findings consistent with encephalitis were observed in 10/18 (Table 1). Most patients had fever (13/18) and lethargy/drowsiness (11/18). Irritability was recorded in 6/18 patients, ataxia/unusual behaviour in 7/18, decreased level of consciousness (LOC) in 5/18, confusion in 4/18, headache in 6/18, vomiting in 5/18, poor feeding in 4/18 and status epilepticus in 2/18 (Table 1). In addition, diarrhoea, cough, rash, convergent squint, coryza, eye deviation, dysarthria, facial droop, urinary retention and hyperreflexia were observed in one or two cases (Table 1). One patient was reported with respiratory illness in the two weeks prior to admission. Three patients were transferred to an intensive care unit (ICU) within 24 hours of admission. One patient with severe combined immunodeficiency (SCID) died. Two patients showed clinical improvement with corticosteroid treatment.

Diagnostic testing for a range of viral agents by PCR on CSF, blood/plasma, sputum and stool samples were performed in the local hospitals. All were negative with the exception of one positive detection of rotavirus from a stool sample, and one detection each of rhinovirus and coronavirus (OC43) in respiratory samples (Table S1). None were considered significant to the clinical presentation. Similarly, serological tests were negative (not consistent with acute infection) for all cases (Table S1). Some cases were also tested for antibodies against ganglioside, muscle specific tyrosine kinase (MUSK), acetylcholine receptor (AChR), N-Methyl-D-Aspartate receptor (NMDAR), voltage-gated potassium channel (VGKC) and neuromyelitis optica (NMO). All were negative (Table S1).

A multidisciplinary expert team (PNB, RD, AK, CAJ) re-reviewed clinical presentation, available diagnostic testing using published criteria for assigning causation in encephalitis and criteria for clinically diagnosing autoimmune encephalitis [16, 17]. Following this review, nine were considered to likely have infectious causes, six immune-mediated causes, and three could not be further classified (Table 2).

**Table 2.**
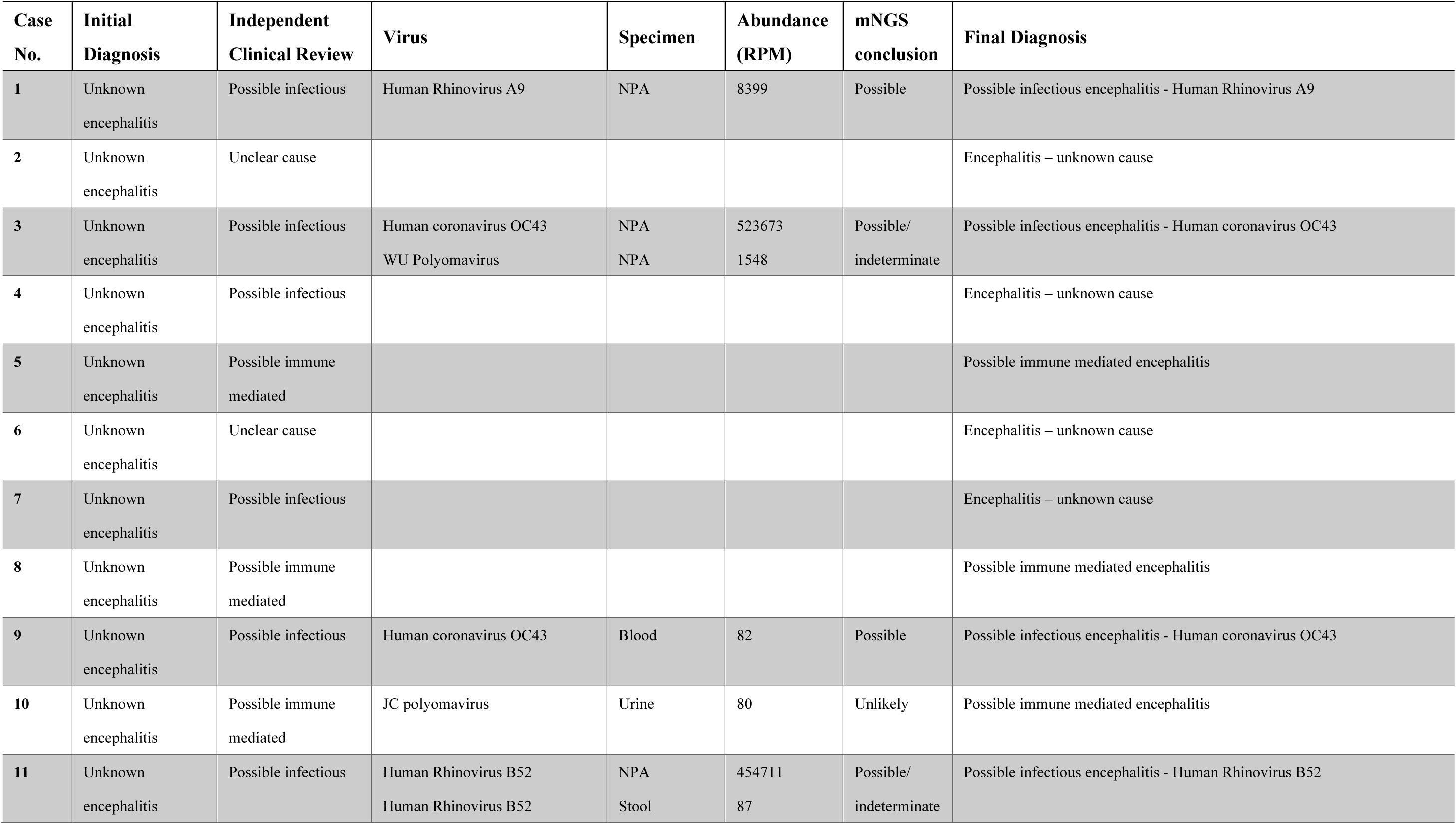
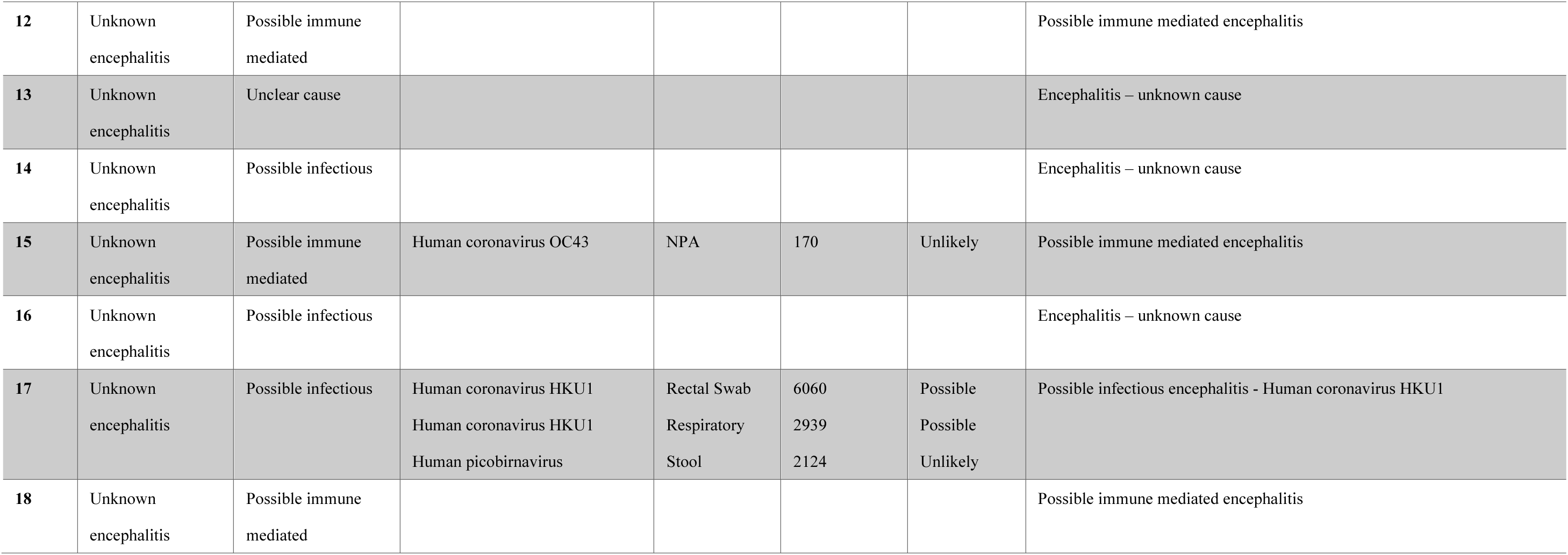
Possible pathogen identification by metagenomic next-generation sequencing (mNGS) and final diagnosis.

| Case No. | Initial Diagnosis | Independent Clinical Review | Virus | Specimen | Abundance (RPM) | mNGS conclusion | Final Diagnosis |
| --- | --- | --- | --- | --- | --- | --- | --- |
| 1 | Unknown encephalitis | Possible infectious | Human Rhinovirus A9 | NPA | 8399 | Possible | Possible infectious encephalitis - Human Rhinovirus A9 |
| 2 | Unknown encephalitis | Unclear cause |  |  |  |  | Encephalitis – unknown cause |
| 3 | Unknown encephalitis | Possible infectious | Human coronavirus OC43<br>WU Polyomavirus | NPA<br>NPA | 523673<br>1548 | Possible/<br>indeterminate | Possible infectious encephalitis - Human coronavirus OC43 |
| 4 | Unknown encephalitis | Possible infectious |  |  |  |  | Encephalitis – unknown cause |
| 5 | Unknown encephalitis | Possible immune mediated |  |  |  |  | Possible immune mediated encephalitis |
| 6 | Unknown encephalitis | Unclear cause |  |  |  |  | Encephalitis – unknown cause |
| 7 | Unknown encephalitis | Possible infectious |  |  |  |  | Encephalitis – unknown cause |
| 8 | Unknown encephalitis | Possible immune mediated |  |  |  |  | Possible immune mediated encephalitis |
| 9 | Unknown encephalitis | Possible infectious | Human coronavirus OC43 | Blood | 82 | Possible | Possible infectious encephalitis - Human coronavirus OC43 |
| 10 | Unknown encephalitis | Possible immune mediated | JC polyomavirus | Urine | 80 | Unlikely | Possible immune mediated encephalitis |
| 11 | Unknown encephalitis | Possible infectious | Human Rhinovirus B52<br>Human Rhinovirus B52 | NPA<br>Stool | 454711<br>87 | Possible/<br>indeterminate | Possible infectious encephalitis - Human Rhinovirus B52 |
| 12 | Unknown encephalitis | Possible immune mediated |  |  |  |  | Possible immune mediated encephalitis |
| 13 | Unknown encephalitis | Unclear cause |  |  |  |  | Encephalitis – unknown cause |
| 14 | Unknown encephalitis | Possible infectious |  |  |  |  | Encephalitis – unknown cause |
| 15 | Unknown encephalitis | Possible immune mediated | Human coronavirus OC43 | NPA | 170 | Unlikely | Possible immune mediated encephalitis |
| 16 | Unknown encephalitis | Possible infectious |  |  |  |  | Encephalitis – unknown cause |
| 17 | Unknown encephalitis | Possible infectious | Human coronavirus HKU1<br>Human coronavirus HKU1<br>Human picobirnavirus | Rectal Swab<br>Respiratory<br>Stool | 6060<br>2939<br>2124 | Possible<br>Possible<br>Unlikely | Possible infectious encephalitis - Human coronavirus HKU1 |
| 18 | Unknown encephalitis | Possible immune mediated |  |  |  |  | Possible immune mediated encephalitis |

### Meta-transcriptomic exploration of potential infectious microbes

A total of 43 specimens, comprising CSF, blood, serum, urine, stool, rectal swabs (rectal SW), nasal swabs (respiratory SW), throat swabs (throat SW), nasopharyngeal aspirates (NPA) and endotracheal aspirates (ETA), were utilised in metagenomic testing (Figure 1B). All 43 samples were individually examined using meta-transcriptomics, generating 2.77 billion raw paired-end reads in total (between 12.2 and 81.0 million reads per sample) (Table S2). For each library, 48.46 to 85.79% of the reads were retained after removal of low complexity and redundant reads (Figure S1, Table S2), and 1.10 to 80.19% of these reads were subsequently retained after removal of human reads (Figure S1, Table S2). The resultant sequence reads and assembled contigs were annotated against NCBI reference databases, revealing a number of microbes, including potential pathogens. These are described in more detail below.

### Detection of viral sequences in clinical samples of encephalitis

Blastx comparisons against the nr database identified at least 7 virus species (5 RNA viruses and 2 DNA viruses) associated with human infection (Figure 2). The total virus positive rate of specimens was 23% (10/43), while the total virus positive rate of patients was 39% (7/18). All the viruses identified are known to be associated with overt disease, with the exception of a picobirnavirus present in one stool sample that may represent a bacteriophage [18]. Among these viruses, human coronavirus OC43 (HCoV-OC43) was present in three cases, while human coronavirus HKU1 (HCoV-HKU1), human rhinovirus A (HRVA), human rhinovirus B (HRVB), JC polyomavirus (JCPyV) and WU polyomavirus (WUPyV) were detected in one case each. Finally, HCoV-HKU1 was detected in both respiratory swab and rectal swab of case 17, while HRVB was detected in both NPA and stool of case 11 (Figure 2). CSF was available for testing from 9/18 cases and we observed a relative absence of possible virus sequence detections amongst these CSF samples.

**Fig. 2.**
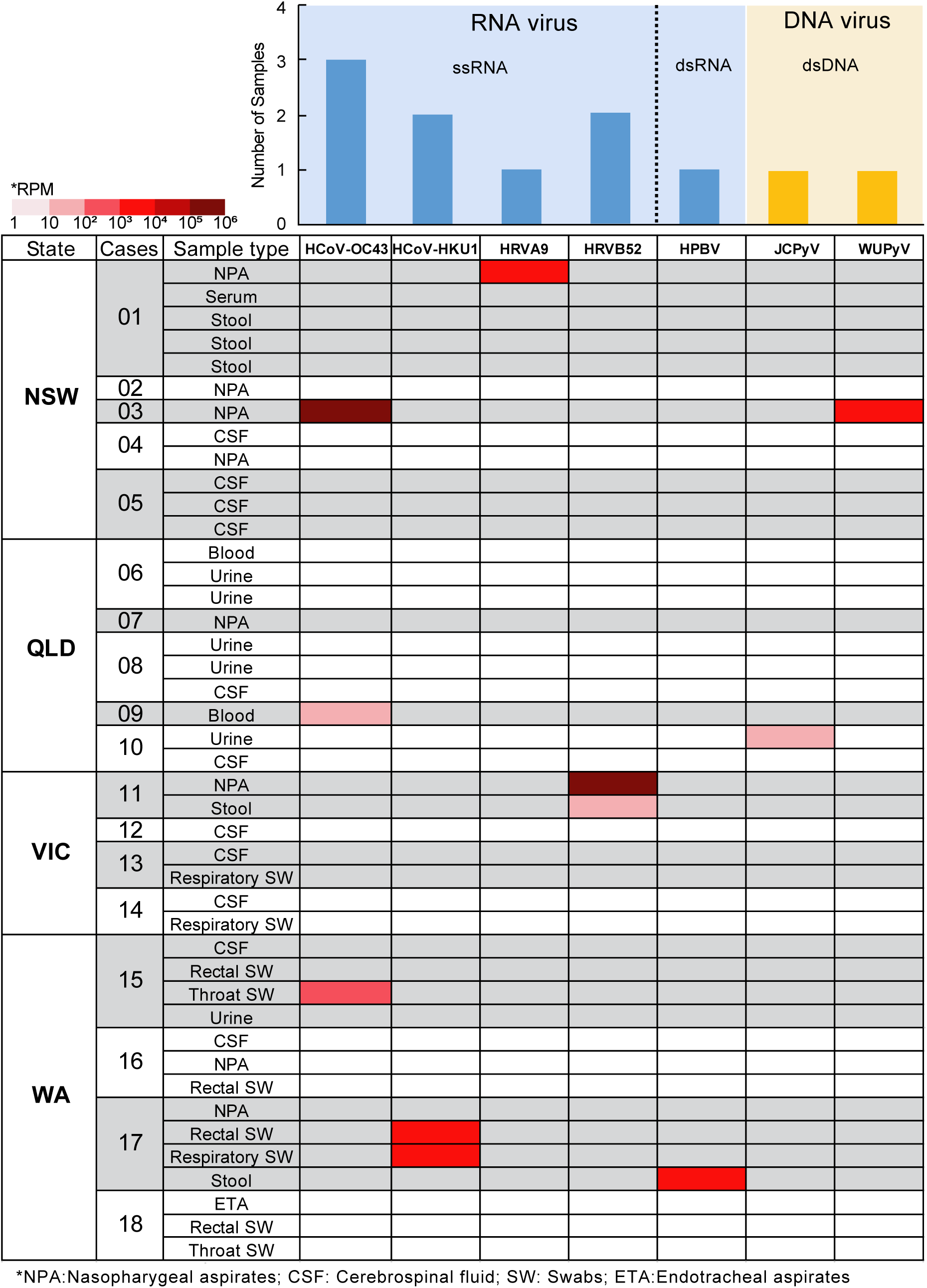
Diversity and abundance of viruses identified in this study. The bar chart shows number of virus species in each sample: RNA viruses (blue) and DNA viruses (yellow). The heatmap shows the abundance level of different virus species within each library. The abundance level of reads was normalized to unique reads mapped per million input reads (RPM). HCoV-OC43, human coronavirus OC43; HCoV-HKU1, human coronavirus HKU1; HRV-A9, human rhinovirus A9; HRV-B52, human rhinovirus B52; Echo6, echovirus E6; HCoV-HKU1, human coronavirus HKU1; HPBV, human picobirnavirus; JCPyV, JC polyomavirus; WUPyV, WU polyomavirus.

Virus expression levels were quantified by estimating their relative abundance (RPM, reads per million). Accordingly, the highest virus abundance was 523,673 RPM (HCoV-OC43, case 03 in NPA), followed by 454,710 RPM (HRV-B52, case 11 in NPA) (Figure 2, Table 2). Viruses with greater than 1,000 RPM (> 0.1% of total reads) included HCoV-HKU1 (case 17, in rectal swab and respiratory swab), HRV-A9 (case 01, in NPA), HPBV (case 17, in stool) and WU polyomavirus (case 03, in NPA) (Figure 2, Table 2). The abundance level of HCoV OC43-was greater than 100 RPM in case 15 (throat swab) and lower than 100 RPM in case 09 (blood) (Figure 2, Table 2).

To identify specific viral genotypes/lineages and their epidemiological origins, phylogenetic analyses were performed using the resulting complete or partial virus genomes (Table S3). This revealed that all HCoV-HKU1, HCoV-OC43, WUPyV and JCPyV sequences determined here were >99.5% identical to the most closely related sequences available on publicly available data bases (Table S3), belonging to genotypes B, G, 3b and 2B, respectively (Figure3A, 3B, 3E and 3F). Two sequences of human rhinovirus B52 - HRV-B52/11-4818/VIC/AU/2019 and HRV-B52/11-4817/VIC/AU/2019 - detected in an NPA and stool specimen from case 11 shared 100% sequence identity with each other yet were relatively distant to known viruses (92.8% nt identity) (Figure 3C, Table S3), such that it likely represents a new genotype of this virus. The human rhinovirus A9 identified here, HRV-A9/01-14618/NSW/AU/2019, exhibited 96.5% sequence identity to the most closely publicly available sequence (Figure 3D, Table S3).

**Fig. 3.**
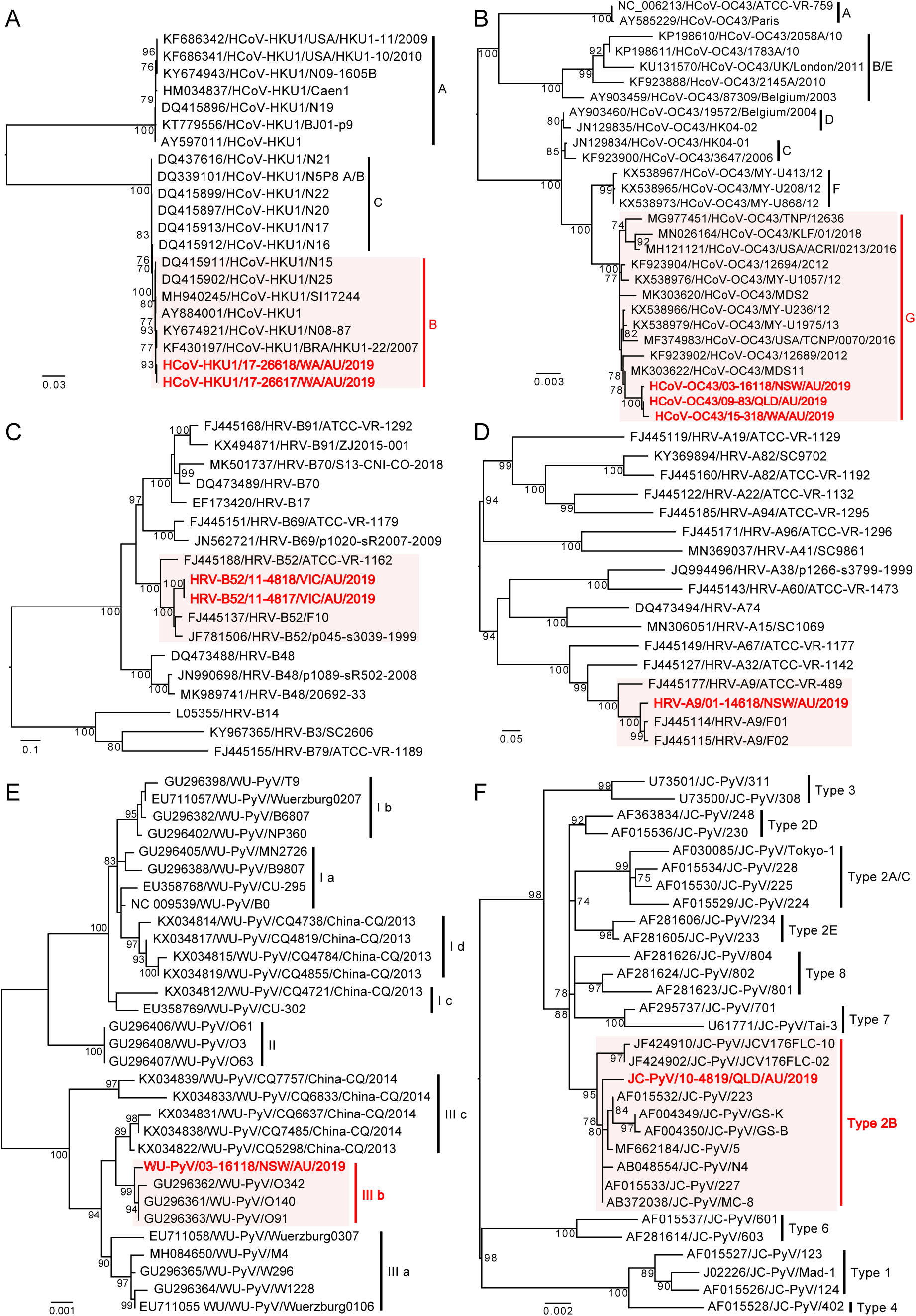
Phylogenetic relationships and intra-specific diversity and of viruses identified by mNGS in this study. Viruses identified as part of this project are marked in red and viruses from the same type/subgroup are shaded light red, whereas those representing background phylogenetic diversity are shown in black. All horizontal branch lengths are scaled to the number of nucleotide substitutions per site, and trees are mid-point rooted for clarity.

One picobirnavirus was identified from one stool sample (case 17), and the near complete sequences of two segments were obtained (Figure 4B). Segment 1 contains two open reading frames, with ORF2 encoding the viral capsid protein and ORF1 encoding a hypothetical protein with unknown function, while segment 2 contains an ORF encoding the viral RNA-dependent RNA polymerase (RdRp) gene (Figure 4B). The conserved motif of the ribosomal binding site (RBS) nucleotide sequence (AGGAGG) is present upstream of the ORF2 of segment 1 and RdRp of segment 2 (Figure 4B). Two copies of the conserved ExxRxxNxxxE aa motif are identifiable un ORF2 of segment 1(Figure 4B). Phylogenetic analysis based on RdRp showed that HPBV/17-26618/WA/AU/2019 is related to primate PBVs within genogroup 1 (Figure 4A), and the RdRp shared 72.4% amino acid sequence identity with the corresponding protein of macaque PBV (AVD54068) (Table S3).

**Fig. 4.**
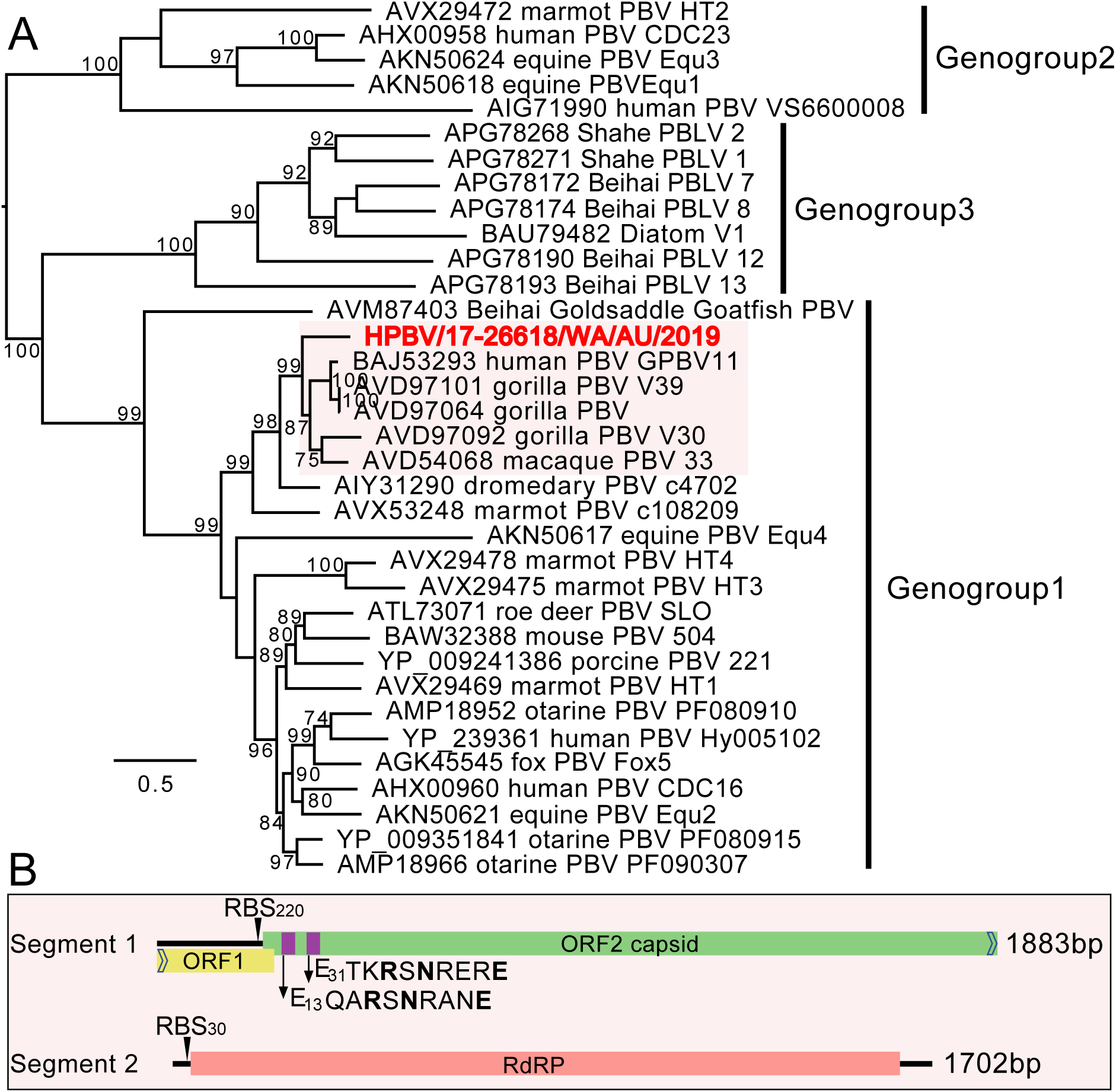
Analysis of the novel picobirnavirus. (A) Phylogenetic relationships of the RNA-dependent RNA polymerase (RdRp) proteins of the novel human picobirnavirus identified from this study and representative picobirnaviruses. (B) Genome organization of the novel human picobirnavirus from one stool sample. Segment 1 encodes two hypothetical proteins (ORF1 and ORF2), with arrows standing for directions of uncomplete ORFs. Segment 2 encodes the viral RNA-dependent RNA polymerase (RdRp). The ribosome binding sites (RBS) and the ExxRxNxxxE amino acid (aa) motifs are marked.

### Characterization of infecting bacteria

We characterised microbial taxonomic clades belonging to seven phyla and 74 species (Figure 5A). The dominant bacterial families identified were *Enterobacteriaceae* (63.64%, estimated using MetaPhlAn2), followed by *Veillonellacae* (10.18%), *Enterococcaceae* (6.07%), *Pseudomonadaceae* (5.89%), *Clostridiales family xi incertae* (5.30%), *Streptococcaceae* (2.75%), and *Bacteroidaceae* (1.45%) (Figure 5). As *Escherichia coli*, unclassified *Escherichia* and unclassified *Veillonella* were observed in almost every library (Figure 6), they were considered unlikely to be infectious agents. *Enterococcus faecalis*, as part of intestinal microbiota and an opportunistic pathogen, was detected in two sterile sites: blood (case 06) and CSF (case 14), although with extremely low abundance (1 and 14 RPM; 53 and 549 reads, respectively) (Figure 6). In other cases, the bacteria identified were known human colonisers or of indeterminate pathogenic status and were observed in non-sterile sites making them unlikely pathogens (Figure 6). No bacterial sequences detected were considered potentially pathogenic.

**Fig. 5.**
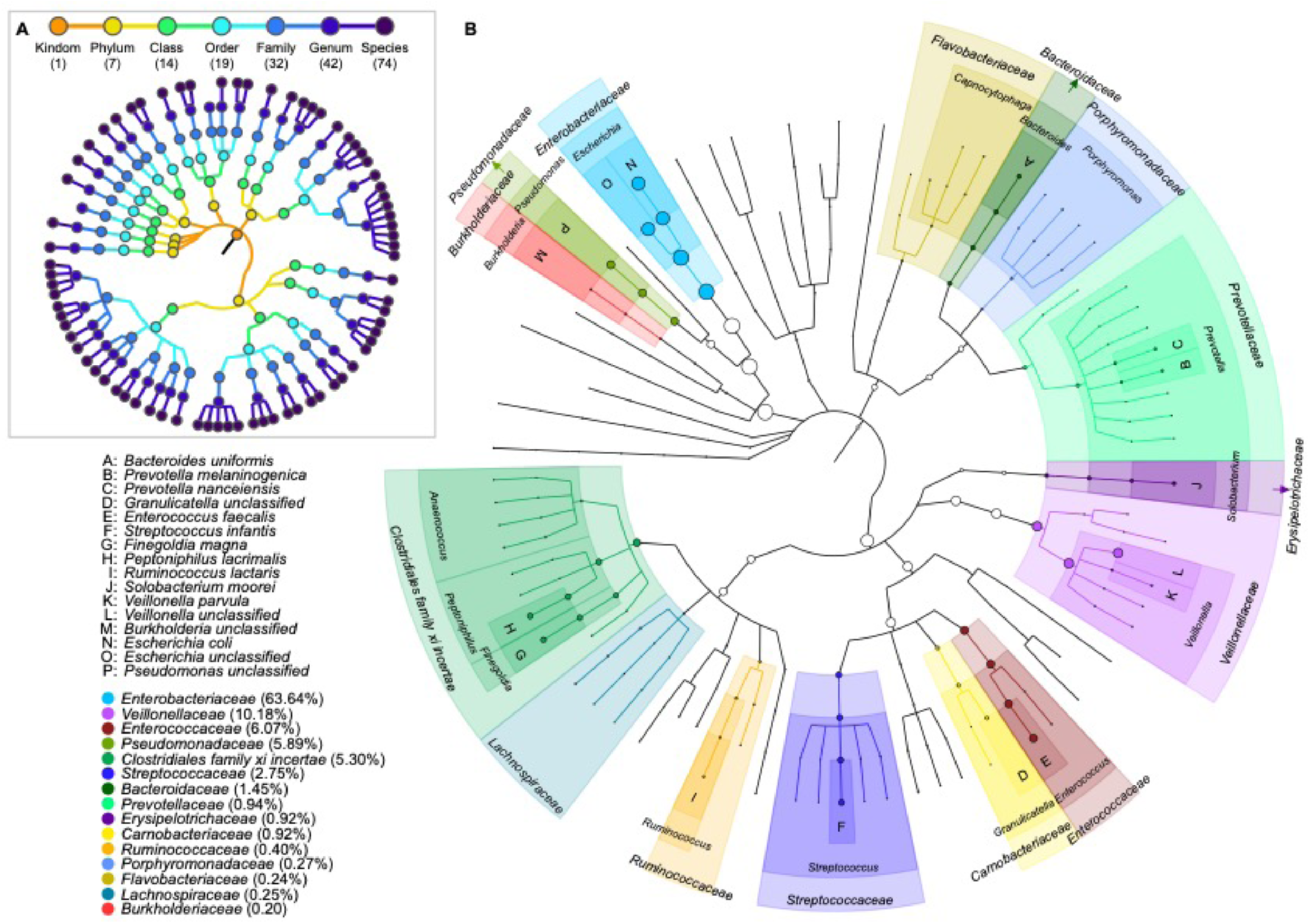
Detailed analysis of the bacteria identified in this study. (A) Bacterial taxa identified using mNGS. Each dot represents a taxonomic entity. From the inner to outer circles, the taxonomic levels range from kingdom to species. Different colored dots indicate different taxonomic levels according to the color key shown. Numbers in parentheses indicate the total number of unique taxonomies detected at each level. (B) Profiling the relative abundance of the bacteria identified here. Cladogram representing the predominant family and top 16 bacterial species present in these cases. Circle size depicts microbial abundance. Abundance levels (reads per million total reads) were estimated using MetaPhlAn2. The taxonomic tree was visualized using GraPhlAn.

**Fig. 6.**
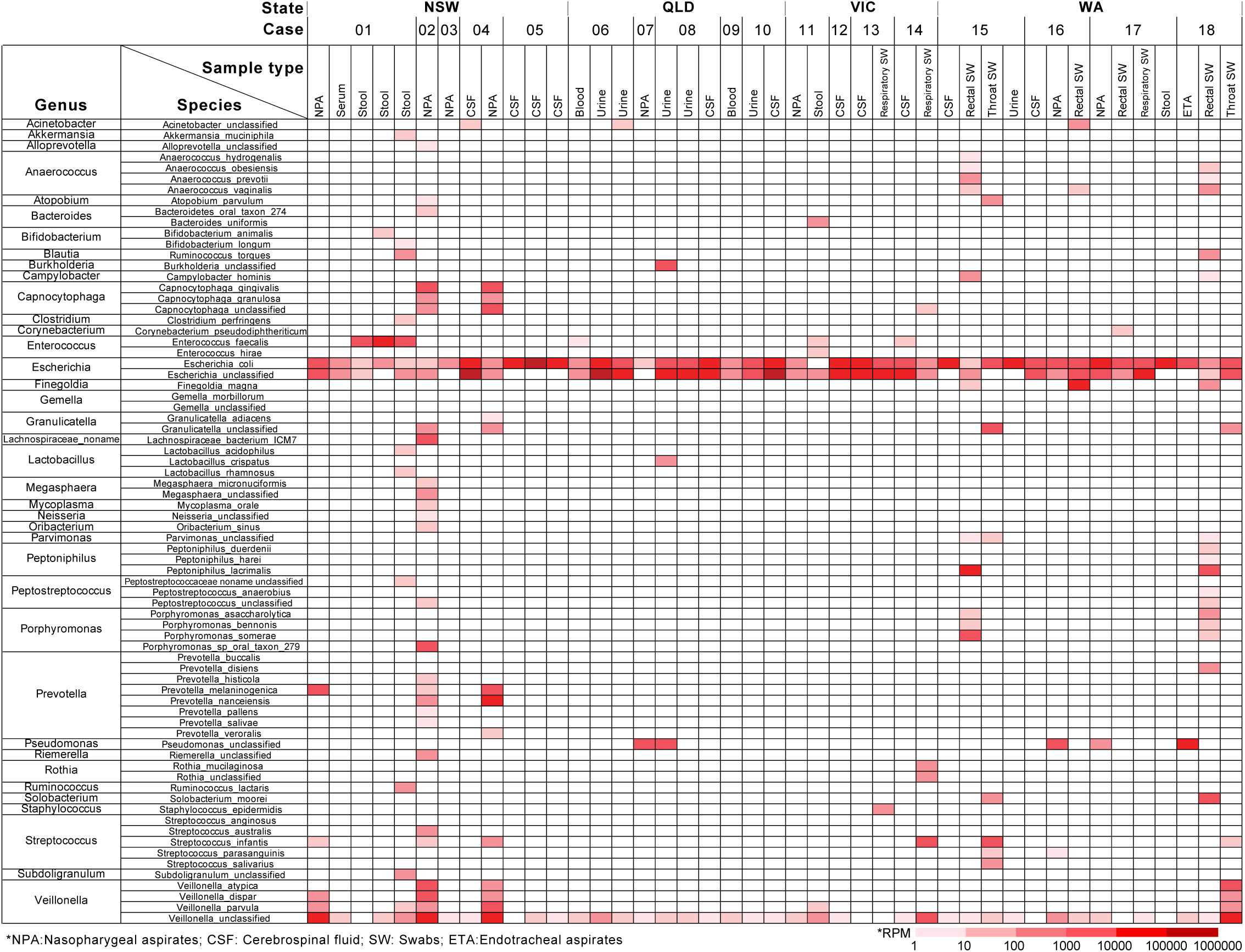
Heatmap of the abundance level for the bacteria identified in each case studied here. The abundance level of microbial reads was normalized to RPM, and the estimation was performed at the species level.

### Correlation between results from clinical evaluation and mNGS

Among those cases with likely infectious causes following clinical evaluation, seven harboured viruses at non-sterile sites (NPA, urine, stool, throat swab, rectal swab and respiratory swab) and one in a sterile site (blood) (Table 2). Case 03, a 1-year-old boy with severe combined immunodeficiency (SCID), had the highest abundance of human coronavirus OC43 (523,673 RPM) and medium abundance of Wu polyomavirus (1548RPM) in NPA, indicating a possible synergistic interaction between these two viruses. Although human coronavirus OC43 has not usually been considered a pathogen capable of causing encephalitis, it’s very high abundance and clinically likely infectious cause makes it a possible pathogen in this case. Human rhinoviruses (HRV-A9 and HRV-B52), that are commonly associated with respiratory diseases, were detected in NPA of case 01 and 11, with high to extremely high abundance (8399-454,711 RPM) (Figure 2, Table 2), respectively. These cases were clinically considered likely infectious, suggesting that the two rhinoviruses identified may be the causative pathogens. JC polyomavirus and human coronavirus OC43 were detected in cases 10 and 15 at relatively low abundance (80 and 170 RPM, respectively). Although both these viruses have previously been associated with encephalitis [19, 20], their low abundance and potential constitutive expression (JCV) make them less likely to be pathogens in these two cases. Importantly, although HCoV-OC43 was not highly abundant in case 09, it was detected in blood (sterile site), which indicates likely pathogenicity in this case (Figure 2, Table 2). In addition, a high abundance of human coronavirus HKU1 were detected in both respiratory swab and rectal swab of case 17, who was reported with a respiratory illness in the before admission, suggesting potential pathogenicity in this case.

Among those cases with likely immune-mediated cause following clinical evaluation, viruses were detected in only two: case 10 (JCV) and case 15 (HCoV-OC43). However, in both cases the viruses were at very low abundance and present at non-sterile sites and so were considered unlikely pathogens. The relative lack of pathogen sequences among those cases with likely immune-mediated causes was notable.

## Discussion

Identification of a causative agent in infectious encephalitis is most validly achieved by obtaining a specimen directly at the site of inflammation, such as CSF or brain biopsy tissue, although the latter is seen as a last resort as sampling comes with associated risks [4]. Therefore, in most cases, CSF is considered the best surrogate specimen for the assessment of neurological disorders [21]. Unlike routine laboratory methods that are limited to the detection of known or related agents, mNGS offers unprecedented opportunities to detect a broad range of microbes including their relative abundance and has recently been employed successfully in detecting a diverse array of potential pathogens not previously associated with CNS disease [22], including astroviruses [3, 23, 24] and arenavirus [25] in cases of encephalitis in immunocompromised and transplant-associated patients. Also of note was that a pegivirus, although considered non-pathogenic, was detected using mNGS in a patient with encephalitis [26], and this technique has recently detected parasitic worms, bacteria, and fungi in CSF specimens from patients with CNS infection of unknown aetiology [5, 11, 27–29].

Importantly, in instances where a CSF sample is unavailable or mNGS testing of CSF fails to detect a pathogen, other sites and samples including respiratory tract (nasal and throat swabs, NPA), gastrointestinal tract (faeces, rectal swab), urine or blood may provide an additional indirect opportunity to test for the presence of potential pathogens that might have transferred in the brain [4]. Further, for some well-established CNS pathogens, CSF testing for viral nucleic acid shows low sensitivity. For example, flaviviruses and enterovirus A71 have been detected in throat swabs, stool or urine from patients with clinical encephalitis, yet were absent in CSF or plasma/serum collected concurrently [30–32]. Among the cases with CSF available for testing (9/18 cases) here, none showed the presence of a possible viral pathogen. In contrast, four respiratory viruses – human rhinovirus A9, human rhinovirus B52, human coronavirus OC43 and human coronavirus HKU1 – were detected in respiratory and gastrointestinal samples (NPA, respiratory swab and rectal swab) from four cases, all of which were in high abundance (>1,000 RPM), indicating that they are experiencing active replication in these patients. Further, these viruses were detected in patients considered likely to have an infectious aetiology in an independent clinical evaluation and so are considered potential pathogens.

Human coronaviruses are respiratory viruses infrequently reported with neuroinvasive properties in both mice and humans [20, 33, 34]. One study suggested that coronavirus infection in the central nervous system is as common as in respiratory tract, although with distinct features [35]. HCoV-OC43 has been identified in CSF and brain tissue in isolated cases of children with acute encephalomyelitis or ADEM [20, 36]. In mice, HCoV-OC43 has been shown to directly invade the CNS via the olfactory route, indicating that intranasal infection may play a role in propagating viruses [20, 37]. Our detection of HCoV in NPA and respiratory samples in high abundance suggests they are possible causative agents, and that the olfactory pathway needs further consideration as a route, albeit infrequent, for neuroinvasion in humans [33, 38]. The potential role of human coronaviruses in acute neurological disease has been further highlighted by the growing evidence of neurological disease of multiple phenotypes as infrequent clinical presentation of SARS-CoV-2 infection [39–42].

Rhinoviruses (genus *Enterovirus*) are the cause of various respiratory illnesses [38]. Unlike some members within the same genus (polioviruses, echovirus, coxsackieviruses, enteroviruses) often implicated in encephalitis, rhinoviruses are discounted as potential causes of encephalitis in consensus criteria [43]. However, although rhinoviral CNS infection is rare, there are reports of human rhinovirus A and B in respiratory specimens associated with acute neurological disease including acute cerebellitis and meningitis, respectively, and one rhinovirus (unknown species) was detected in CSF in a case of sepsis-like illness [44–46]. Further, enterovirus D68 is acknowledged as an emerging neurotropic enterovirus with predominant replication in the respiratory tract [47, 48], suggesting that members of the *Enterovirus* genus may have neuropathic potential more broadly. Although the two species of rhinovirus detected in this study were identified in in nasopharyngeal aspirates, on the basis of clinical evaluation and mNGS analysis they are considered possible causative agents. Targeted assays of the CSF or antibody tests could be beneficial for pathogen determination in future cases.

Another possible explanation for the pathogenicity of these respiratory viruses at non-CSF sites is that they represent para-infectious encephalitis resulting from indirect CNS pathogenicity. A variety of severe encephalopathy/encephalitis syndromes have been described in association with viruses such as influenza, adenovirus and human herpes virus-6, and it is possible that other respiratory viruses could produce disease via these mechanisms [38, 49–53]. It is clear that larger studies are needed to better understand the role of non-CNS samples in the detection of candidate pathogens amongst cases of encephalitis with otherwise unknown cause. These studies should involve the testing of non-CNS samples in combination with molecular and serological CSF and serum investigations.

The frequency and range of bacterial reads identified in almost all specimens analysed highlights the challenge of applying and interpreting mNGS particularly to non-sterile samples. Most reads corresponded to known human colonisers and/or organisms with questionable pathogenicity. An exception was *E. coli* that was detected in sterile site samples from several cases, but discounted given that reads were present in all samples analysed. Similarly, *E. faecalis* was present in two sterile site samples in two cases, but at very low abundance in sterile site specimens in two cases, and we were unable to confirm these infections by PCR. Notably, all the cases studied received empiric antibiotic treatment with a third-generation cephalosporin to which enterococci are inherently resistant. Concurrent or preceding antibiotic usage is an important consideration in mNGS studies where treatment may bias detection to resistant bacterial species [9].

An additional insight from this study is the apparent utility of clinical definitions of immune-mediated encephalitis. Among the unknown encephalitis cases under enhanced evaluation here, six were considered likely to be immune-mediated. Despite a similar number and spectrum of samples available for mNGS, we found no possible pathogens using mNGS among these six cases compared with possible pathogens in five of nine cases clinically considered likely to be infectious by an independent clinical review panel. The frequency of likely immune-mediated encephalitis in this case series of encephalitis of unknown cause emphasises the need for further research to identify diagnostic biomarkers.

It is not possible to definitively determine that the viruses identified using mNGS here were the causes of the encephalitis in each individual case studied. However, the combination of enhanced clinical evaluation with mNGS appears a fruitful way forward in evaluation of cases of unknown encephalitis [9].

## Methods

### Ethics statement

This study was performed under the ethical approval of the Sydney Children’s Hospitals Network Human Research Ethics Committee (HREC/13/SCHN/191).

### Enrolment criteria and case definition

The cases selected in this study all met the case definition of encephalitis [54]. This included patients aged <= 14 years admitted to hospital with encephalopathy, altered consciousness that persisted for longer than 24 hours, including lethargy, irritability, or a change in personality and behaviour, and at least two of the following criteria: (i) fever or history of fever (≥38°C) during the presenting illness, (ii) seizures and/or focal neurological, (iii) abnormal findings on electroencephalographic (EEG) consistent with encephalitis, and (iv) abnormal results of neuroimaging (CT or MRI) suggestive of encephalitis (Table 1). In addition, each of the study subjects tested negative with routine diagnostic protocols.

All samples were collected from diagnostic laboratories between March 2014 and December 2016. All specimens were originally collected by trained clinicians using sterile swabs or aseptic technique and were immediately sent to local laboratories where diagnostic testing was undertaken. Following diagnostic testing, residual samples were stored at -20°C. Following collection for research purposes, informed consent was obtained from parents/guardians for research testing and specimens were then placed into -80°C storage locally and transferred in batches on dry ice to a central specimen repository where they were stored in a -80°C freezer.

### RNA extraction, library construction and sequencing

Samples were subjected to RNA extraction using the RNeasy plus universal kit (QIAGEN, Chadstone Centre, Victoria, Australia). The concentration and quality of final extractions were examined using a NanoDrop spectrophotometer (ThermoFisher Scientific, USA). The Trio RNA-Seq kit (NuGEN Technologies, USA) was used for library preparation in all cases as it targets low concentration RNA samples (as low as 500 pg) [55]. Paired-end (150 bp) sequencing of these RNA library was performed using the Illumina NovaSeq platform. All library preparation and sequencing steps were performed by Australian Genome Research Facility (AGRF), Melbourne.

### Microbial characterization

For each library/case, quality control was performed and adaptor sequences and low-quality/low-complex reads were removed using BBmap (https://jgi.doe.gov/data-and-tools/bbtools/bb-tools-user-guide/bbmap-guide/) and CD-HIT-DUP [56]. Human reads were removed by mapping to the human genome. Non-human reads were then compared against reference virus database downloaded from GenBank and the NCBI non-redundant protein database using Blastn and Diamond [57] blastx, respectively, with e-value thresholds of 1 × 10^-10^ and 1×10^-4^. Blast hits were then annotated by taxonomy. Reads were *de novo* assembled using Megahit [58, 59] from the virus-positive library in which they were identified based on the blast procedure described above. In cases with low genome coverage, reads were directly mapped to the sequence of a close relative, and a consensus genome was obtained from the mapped reads. To exclude contamination due to index hopping, for each virus only those present at >0.1% of the highest viral abundance were considered true positives. Sequence specific primers were designed, and RT-PCR and Sanger sequencing were performed to verify and confirm the presence of viruses with highly similar sequences that appeared in multiple libraries.

Virus abundance were estimated as Reads Per Million (RPM) [55], calculated by using the relation “mapped reads / total reads * one million”. For picobirnavirus, open reading frames were predicted using orffinder (https://www.ncbi.nlm.nih.gov/orffinder/) and gene annotated against the conserved domain database (CDD) [60].

The resulting complete or partial virus genomes (Table S3), as well as related background viruses from GenBank, were aligned using MAFFT version 7 [61] and subjected to phylogenetic analysis using the maximum likelihood method available in PhyML 3.0 [62] employing the General Time Reversible (GTR) model of nucleotide substitution with a gamma distribution of among-site rate variation and 1000 bootstrap replicates.

Bacterial taxonomic profiling was initially performed using MetaPhlAn2 [63] with default parameters, mapping the non-human sequence reads to a set of ∼1 million unique clade-specific marker genes from ∼13,500 bacterial and archaeal species.

### Data availability

All non-human sequence reads generated here have been deposited on the NCBI Sequence Read Achieve (SRA; BioProject accession PRJNA633210).

### Clinical evaluation

All 18 cases had previously undergone clinical review as part of the Australian Childhood Encephalitis study and been determined to have an unknown cause [7]. A clinical evaluation panel was convened comprising a clinical paediatric neurologist (RD), paediatric infectious disease physician (CJ), clinical epidemiologist (PB) and clinical microbiologist (AK) who re-evaluated each case in terms of clinical presentation, available diagnostic testing and response to empiric therapies using published criteria for assigning causation in encephalitis [43] and clinically diagnosing autoimmune encephalitis [16]. This panel was blinded to the mNGS results.

### Logic model for determining pathogenic potential of mNGS detections

We applied a logic model (Figure S2) to determine the likely significance of mNGS results based upon the nature of the sample from which mNGS detection occurred, the organism abundance and pathogenicity, and accounting for the enhanced clinical evaluation findings to arrive at a final clinical diagnosis.

## Acknowledgments

We acknowledge The University of Sydney HPC service (Artemis) for providing high-performance computing resources that have contributed to the research reported in this paper.

## Author Contributions

**Conceptualization:** Cheryl A. Jones, Philip N. Britton, Edward C. Holmes.

**Data curation:** Ci-Xiu Li, Rebecca Burrell.

**Formal analysis:** Ci-Xiu Li, Rebecca Burrell, Philip N. Britton, Edward C. Holmes.

**Funding acquisition:** Cheryl A. Jones, Philip N. Britton, Edward C. Holmes.

**Investigation:** Russell C. Dale, Alison Kesson, Christopher C Blyth, Julia E Clark, Nigel Crawford, Cheryl A. Jones

**Methodology:** Ci-Xiu Li, Rebecca Burrell

**Resources:** Rebecca Burrell, Alison Kesson, Christopher Blyth, Julia Clark, Nigel Crawford.

**Writing - original draft:** Ci-Xiu Li.

**Writing - review & editing:** Ci-Xiu Li, Rebecca Burrell, Russell C. Dale, Christopher C Blyth Julia E Clark, Cheryl A. Jones, Philip N. Britton, Edward C. Holmes.

## Supporting Information Captions

**Fig S1.**
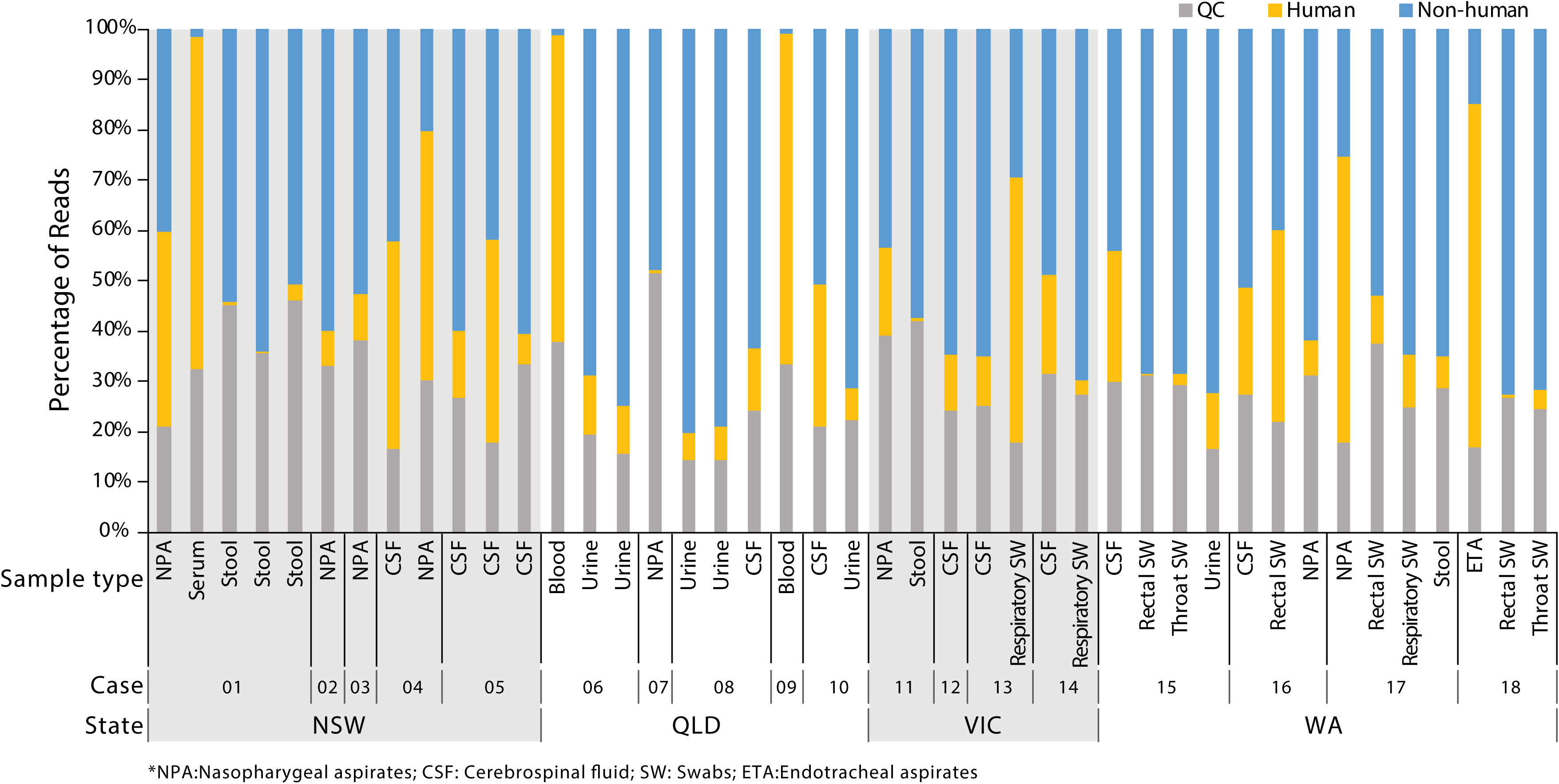
Summary of taxonomy and abundance from meta-transcriptomic sequencing of the libraries from each patient. The bar chart shows the proportion of assigned taxa according to the keys provided. The panel shows the proportion of reads that were assigned as quality controlled (QC), human and non-human in each library.

**Fig S2.**
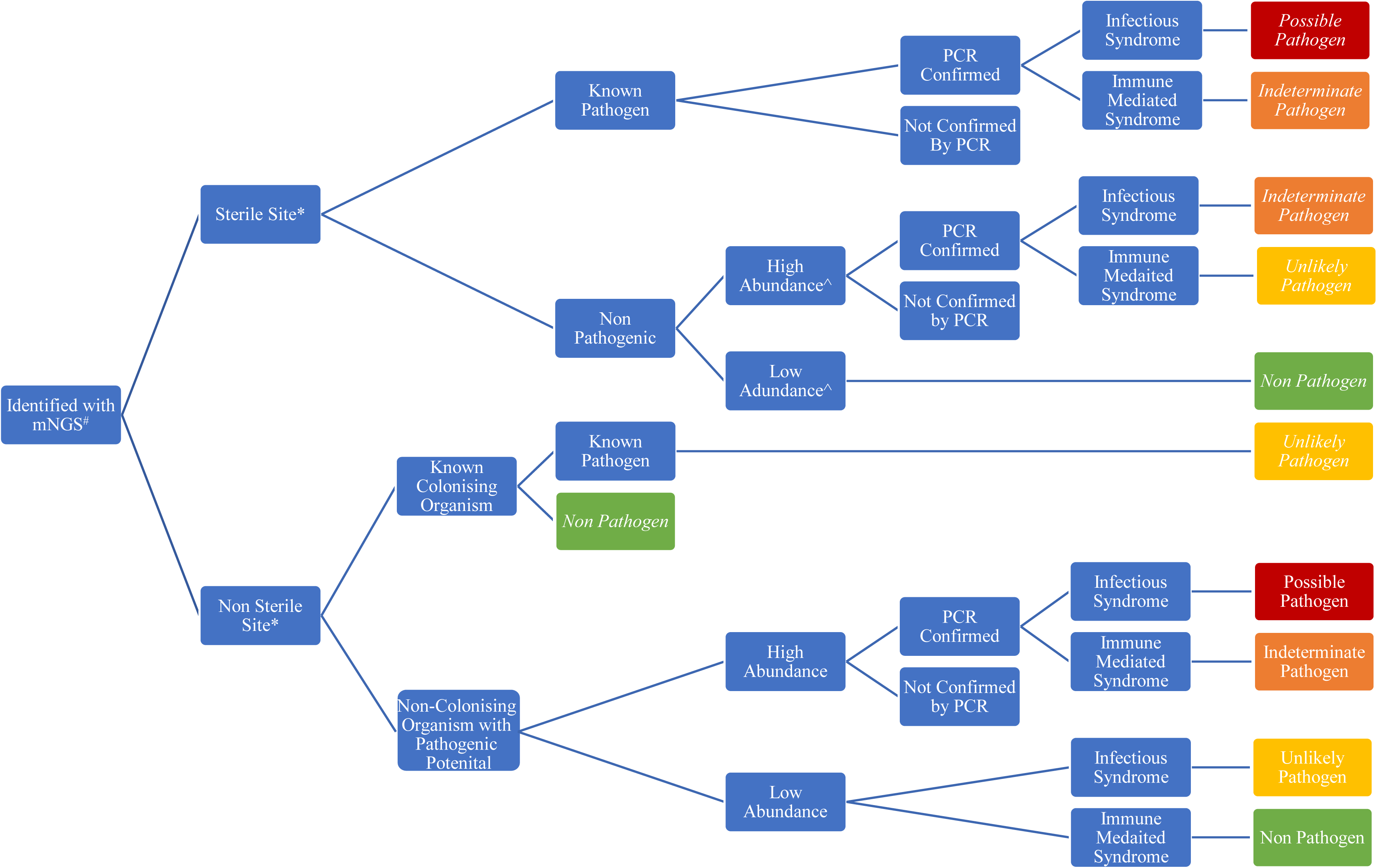
Logic model to determine potential the pathogenicity of the viruses, bacteria and fungi identified here. #mNGS = metagenomic next-generation sequencing. *Sterile sites included blood, serum, and CSF; non-sterile sites included nasopharyngeal aspirates, endotracheal samples, throat and nasal swabs, as well as stool and rectal swabs. Infectious and immune-mediated syndromes were defined by the advanced clinical expert panel. ^Abundance was determined to be high if sequence reads were above 10000 RPM, or low if below 1000 RPM. In the event of detection in multiple samples sterile samples taken priority.

**Table S1.**
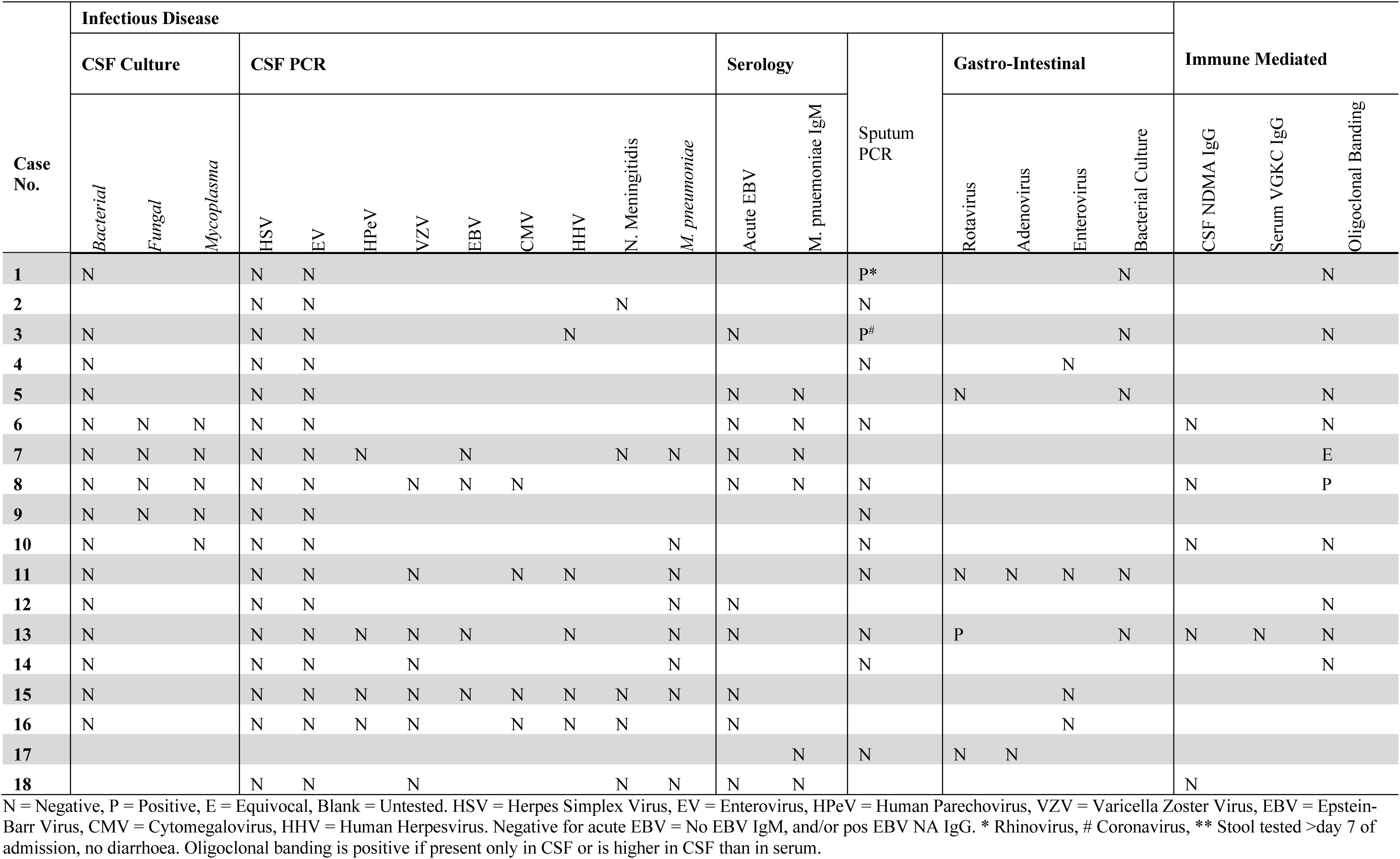
Summary of laboratory tests performed on the encephalitis cases.

**Table S2.**
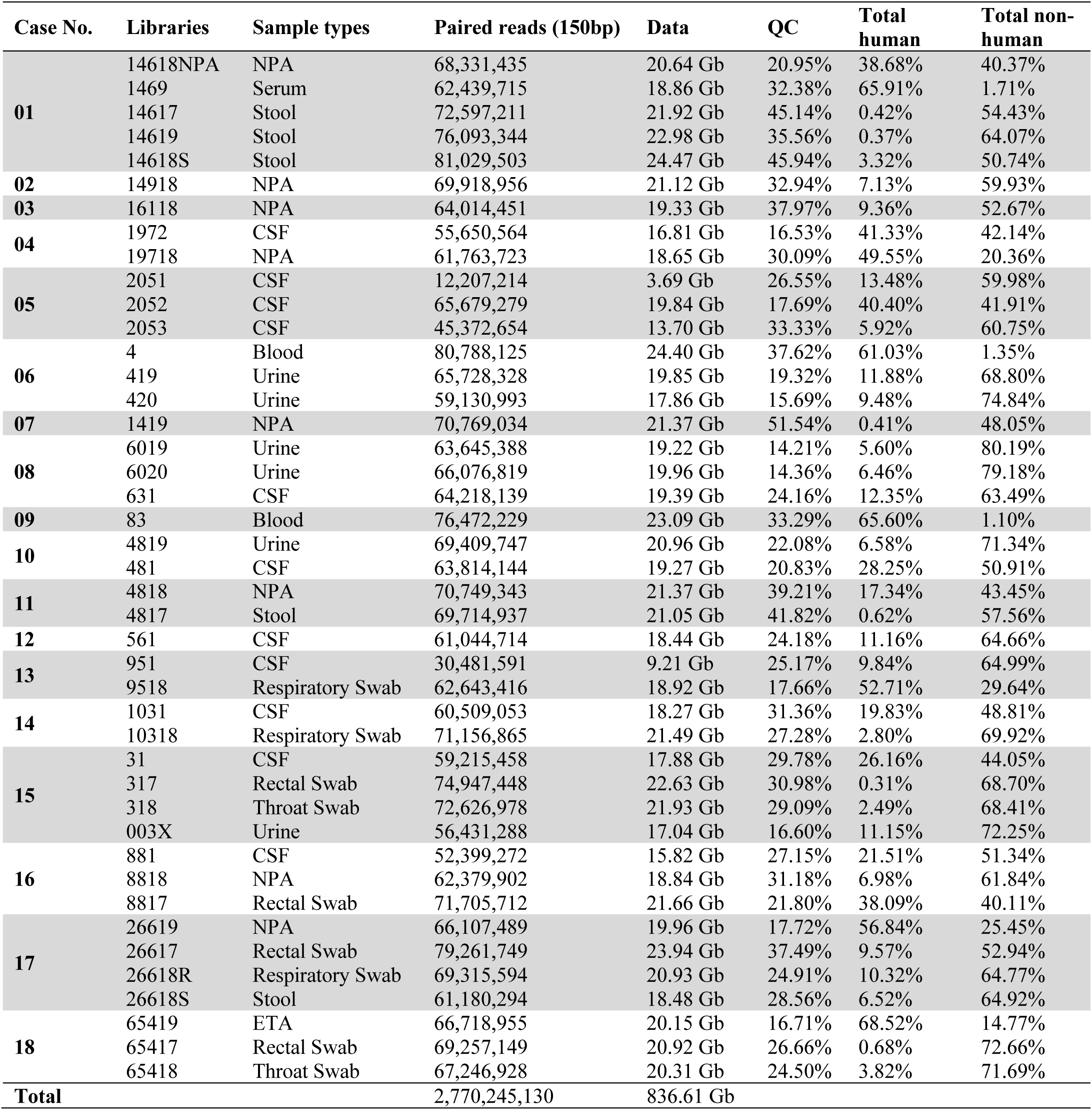
Information on the RNA sequencing libraries generated in this study.

**Table S3.**
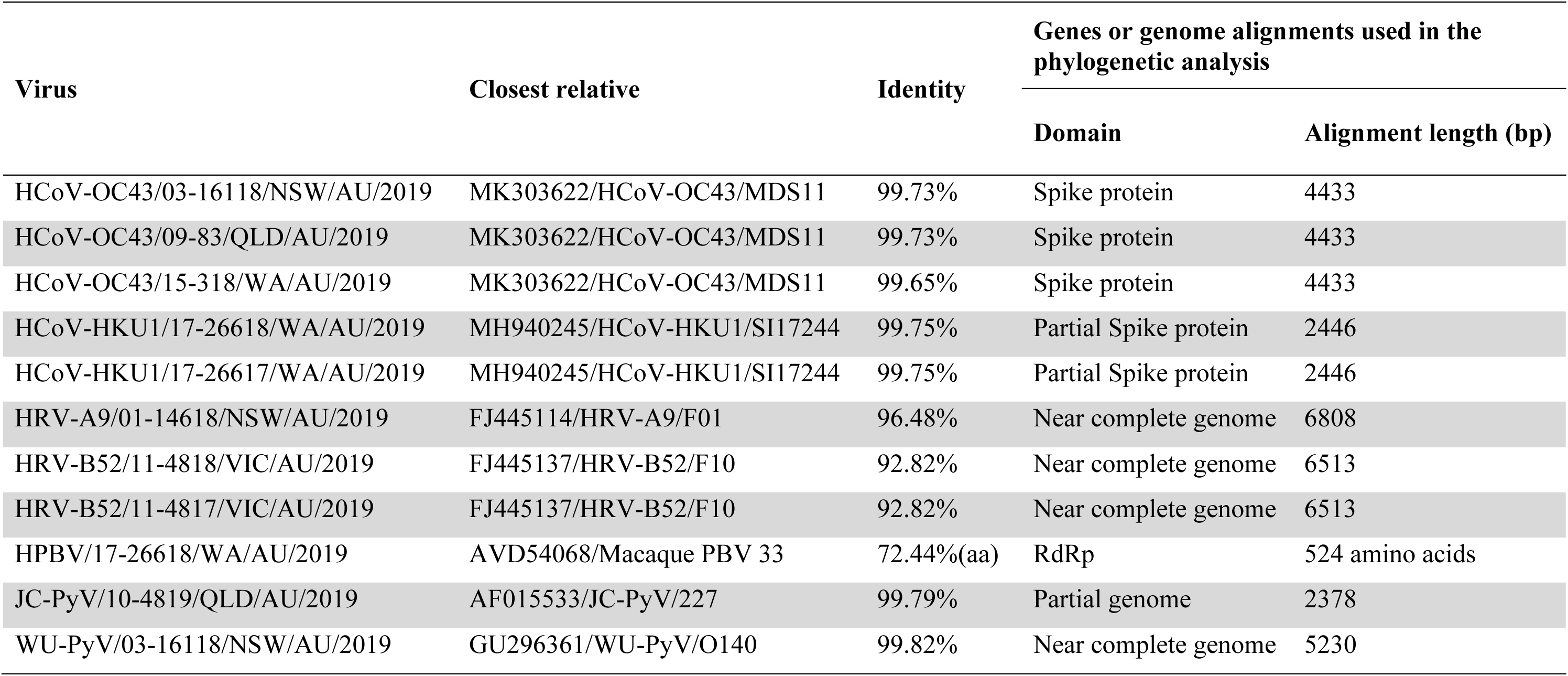
Identity of viruses identified in this study with the most closely related sequence available on public sequence databases.

