## Supplementary Information for "Diagnosis and analysis of unexplained cases of childhood encephalitis in Australia using metagenomic next-generation sequencing"

**Table S1.** Summary of laboratory tests performed on the encephalitis cases.

| Case No. | **Infectious Disease** | | | | | | | | | | | | | | | | | | | | **Immune Mediated** | |
|---|---|---|---|---|---|---|---|---|---|---|---|---|---|---|---|---|---|---|---|---|---|---|
| | **CSF Culture** | | | **CSF PCR** | | | | | | | | | **Serology** | | Sputum PCR | **Gastro-Intestinal** | | | | | | |
| | *Bacterial* | *Fungal* | *Mycoplasma* | HSV | EV | HPeV | VZV | EBV | CMV | HHV | N. Meningitidis | *M. pneumoniae* | Acute EBV | M. pnuemoniae IgM | | Rotavirus | Adenovirus | Enterovirus | Bacterial Culture | CSF NDMA IgG | Serum VGKC IgG | Oligoclonal Banding |
| **1** | N | | | N | N | | | | | | | | | | P* | | | | N | | | N |
| **2** | | | | N | N | | | | | | N | | | | N | | | | | | | |
| **3** | N | | | N | N | | | | | N | | | N | | P# | | | | N | | | N |
| **4** | N | | | N | N | | | | | | | | | | N | | | N | | | | |
| **5** | N | | | N | N | | | | | | | | N | N | | N | | | N | | | N |
| **6** | N | N | N | N | N | | | | | | | | N | N | N | | | | | N | | N |
| **7** | N | N | N | N | N | N | | N | | | N | N | N | N | | | | | | | | E |
| **8** | N | N | N | N | N | | N | N | N | | | | N | N | N | | | | | N | | P |
| **9** | N | N | N | N | N | | | | | | | | | | N | | | | | | | |
| **10** | N | | N | N | N | | | | | | | N | | | N | | | | | N | | N |
| **11** | N | | | N | N | | N | | N | N | | N | | | N | N | N | N | N | | | |
| **12** | N | | | N | N | | | | | | | N | N | | | | | | | | | N |
| **13** | N | | | N | N | N | N | N | | N | | N | N | | N | P | | | N | N | N | N |
| **14** | N | | | N | N | | N | | | | | N | | | N | | | | | | | N |
| **15** | N | | | N | N | N | N | N | N | N | N | N | N | | | | | N | | | | |
| **16** | N | | | N | N | N | N | | N | N | N | | N | | | | | N | | | | |
| **17** | | | | | | | | | | | | | | N | N | N | N | | | | | |
| **18** | | | | N | N | | N | | | | N | N | N | N | | | | | | N | | |

N = Negative, P = Positive, E = Equivocal, Blank = Untested. HSV = Herpes Simplex Virus, EV = Enterovirus, HPeV = Human Parechovirus, VZV = Varicella Zoster Virus, EBV = Epstein-Barr Virus, CMV = Cytomegalovirus, HHV = Human Herpesvirus. Negative for acute EBV = No EBV IgM, and/or pos EBV NA IgG. * Rhinovirus, # Coronavirus, ** Stool tested >day 7 of admission, no diarrhoea. Oligoclonal banding is positive if present only in CSF or is higher in CSF than in serum.

**Table S2**. Information on the RNA sequencing libraries generated in this study.

| Case No. | Libraries | Sample types | Paired reads (150bp) | Data | QC | Total human | Total non-human |
|---|---|---|---|---|---|---|---|
| **01** | 14618NPA | NPA | 68,331,435 | 20.64 Gb | 20.95% | 38.68% | 40.37% |
| | 1469 | Serum | 62,439,715 | 18.86 Gb | 32.38% | 65.91% | 1.71% |
| | 14617 | Stool | 72,597,211 | 21.92 Gb | 45.14% | 0.42% | 54.43% |
| | 14619 | Stool | 76,093,344 | 22.98 Gb | 35.56% | 0.37% | 64.07% |
| | 14618S | Stool | 81,029,503 | 24.47 Gb | 45.94% | 3.32% | 50.74% |
| **02** | 14918 | NPA | 69,918,956 | 21.12 Gb | 32.94% | 7.13% | 59.93% |
| **03** | 16118 | NPA | 64,014,451 | 19.33 Gb | 37.97% | 9.36% | 52.67% |
| **04** | 1972 | CSF | 55,650,564 | 16.81 Gb | 16.53% | 41.33% | 42.14% |
| | 19718 | NPA | 61,763,723 | 18.65 Gb | 30.09% | 49.55% | 20.36% |
| **05** | 2051 | CSF | 12,207,214 | 3.69 Gb | 26.55% | 13.48% | 59.98% |
| | 2052 | CSF | 65,679,279 | 19.84 Gb | 17.69% | 40.40% | 41.91% |
| | 2053 | CSF | 45,372,654 | 13.70 Gb | 33.33% | 5.92% | 60.75% |
| **06** | 4 | Blood | 80,788,125 | 24.40 Gb | 37.62% | 61.03% | 1.35% |
| | 419 | Urine | 65,728,328 | 19.85 Gb | 19.32% | 11.88% | 68.80% |
| | 420 | Urine | 59,130,993 | 17.86 Gb | 15.69% | 9.48% | 74.84% |
| **07** | 1419 | NPA | 70,769,034 | 21.37 Gb | 51.54% | 0.41% | 48.05% |
| **08** | 6019 | Urine | 63,645,388 | 19.22 Gb | 14.21% | 5.60% | 80.19% |
| | 6020 | Urine | 66,076,819 | 19.96 Gb | 14.36% | 6.46% | 79.18% |
| | 631 | CSF | 64,218,139 | 19.39 Gb | 24.16% | 12.35% | 63.49% |
| **09** | 83 | Blood | 76,472,229 | 23.09 Gb | 33.29% | 65.60% | 1.10% |
| **10** | 4819 | Urine | 69,409,747 | 20.96 Gb | 22.08% | 6.58% | 71.34% |
| | 481 | CSF | 63,814,144 | 19.27 Gb | 20.83% | 28.25% | 50.91% |
| **11** | 4818 | NPA | 70,749,343 | 21.37 Gb | 39.21% | 17.34% | 43.45% |
| | 4817 | Stool | 69,714,937 | 21.05 Gb | 41.82% | 0.62% | 57.56% |
| **12** | 561 | CSF | 61,044,714 | 18.44 Gb | 24.18% | 11.16% | 64.66% |
| **13** | 951 | CSF | 30,481,591 | 9.21 Gb | 25.17% | 9.84% | 64.99% |
| | 9518 | Respiratory Swab | 62,643,416 | 18.92 Gb | 17.66% | 52.71% | 29.64% |
| **14** | 1031 | CSF | 60,509,053 | 18.27 Gb | 31.36% | 19.83% | 48.81% |
| | 10318 | Respiratory Swab | 71,156,865 | 21.49 Gb | 27.28% | 2.80% | 69.92% |
| **15** | 31 | CSF | 59,215,458 | 17.88 Gb | 29.78% | 26.16% | 44.05% |
| | 317 | Rectal Swab | 74,947,448 | 22.63 Gb | 30.98% | 0.31% | 68.70% |
| | 318 | Throat Swab | 72,626,978 | 21.93 Gb | 29.09% | 2.49% | 68.41% |
| | 003X | Urine | 56,431,288 | 17.04 Gb | 16.60% | 11.15% | 72.25% |
| **16** | 881 | CSF | 52,399,272 | 15.82 Gb | 27.15% | 21.51% | 51.34% |
| | 8818 | NPA | 62,379,902 | 18.84 Gb | 31.18% | 6.98% | 61.84% |
| | 8817 | Rectal Swab | 71,705,712 | 21.66 Gb | 21.80% | 38.09% | 40.11% |
| **17** | 26619 | NPA | 66,107,489 | 19.96 Gb | 17.72% | 56.84% | 25.45% |
| | 26617 | Rectal Swab | 79,261,749 | 23.94 Gb | 37.49% | 9.57% | 52.94% |
| | 26618R | Respiratory Swab | 69,315,594 | 20.93 Gb | 24.91% | 10.32% | 64.77% |
| | 26618S | Stool | 61,180,294 | 18.48 Gb | 28.56% | 6.52% | 64.92% |
| **18** | 65419 | ETA | 66,718,955 | 20.15 Gb | 16.71% | 68.52% | 14.77% |
| | 65417 | Rectal Swab | 69,257,149 | 20.92 Gb | 26.66% | 0.68% | 72.66% |
| | 65418 | Throat Swab | 67,246,928 | 20.31 Gb | 24.50% | 3.82% | 71.69% |
| **Total** | | | 2,770,245,130 | 836.61 Gb | | | |

**Table S3**. Identity of viruses identified in this study with the most closely related sequence available on public sequence databases.

| Virus | Closest relative | Identity | Genes or genome alignments used in the phylogenetic analysis | |
|---|---|---|---|---|
| | | | **Domain** | **Alignment length (bp)** |
| HCoV-OC43/03-16118/NSW/AU/2019 | MK303622/HCoV-OC43/MDS11 | 99.73% | Spike protein | 4433 |
| HCoV-OC43/09-83/QLD/AU/2019 | MK303622/HCoV-OC43/MDS11 | 99.73% | Spike protein | 4433 |
| HCoV-OC43/15-318/WA/AU/2019 | MK303622/HCoV-OC43/MDS11 | 99.65% | Spike protein | 4433 |
| HCoV-HKU1/17-26618/WA/AU/2019 | MH940245/HCoV-HKU1/SI17244 | 99.75% | Partial Spike protein | 2446 |
| HCoV-HKU1/17-26617/WA/AU/2019 | MH940245/HCoV-HKU1/SI17244 | 99.75% | Partial Spike protein | 2446 |
| HRV-A9/01-14618/NSW/AU/2019 | FJ445114/HRV-A9/F01 | 96.48% | Near complete genome | 6808 |
| HRV-B52/11-4818/VIC/AU/2019 | FJ445137/HRV-B52/F10 | 92.82% | Near complete genome | 6513 |
| HRV-B52/11-4817/VIC/AU/2019 | FJ445137/HRV-B52/F10 | 92.82% | Near complete genome | 6513 |
| HPBV/17-26618/WA/AU/2019 | AVD54068/Macaque PBV 33 | 72.44%(aa) | RdRp | 524 amino acids |
| JC-PyV/10-4819/QLD/AU/2019 | AF015533/JC-PyV/227 | 99.79% | Partial genome | 2378 |
| WU-PyV/03-16118/NSW/AU/2019 | GU296361/WU-PyV/O140 | 99.82% | Near complete genome | 5230 |
